# Urine proteome changes in an α-synuclein transgenic mouse model of Parkinson’s disease

**DOI:** 10.1101/2020.04.05.026104

**Authors:** Lujun Li, Xuanzhen Pan, Ting Wang, Yuanrui Hua, Youhe Gao

## Abstract

Urine accommodates more changes than other fluids, and it is a good source in the search for early sensitive biomarkers. The present study collected urine samples from 2-, 4-, 6-, 8- and 10-month-old α-synuclein transgenic mice. Based on data-independent acquisition (DIA) technology, liquid chromatography-tandem mass spectrometry (LC-MS/MS) was used for quantitative analysis. Seventeen human homologous differential proteins were screened and compared with those in the urine of 2-month-old mice, and 9 proteins were related to Parkinson’s disease (PD). Formin-2, Splicing factor 3A subunit 1, and Isopentenyl-diphosphate Delta-isomerase 1 changed continuously in months 6, 8 and 10. These experiments and analyses demonstrated that the urine proteome reflected the development of α-synuclein transgenic mice and provided clues for the early clinical diagnosis of PD.

---

Parkinson’s disease (PD) is a neurodegenerative disease characterized by progressive motor and non-motor disorders [1]. Its main pathological features are the progressive loss of dopaminergic neurons in the dense substantia nigra and the presence of α-synuclein as the main component of eosinophilic inclusion bodies, i.e., Lewy bodies, in the remaining dopaminergic neurons. PD is the second most common neurodegenerative disease, and it affects approximately 1% of people over 65 years of age [2]. With the aging of the population and the prolongation of life expectancy, the incidence rate of PD is expected to double in the next 20 years [3]. PD seriously affects the quality of life of patients, and it results in a huge social and economic burden [4, 5].

The early diagnosis of PD is of great significance to improve the quality of life of patients and delay the development of the disease. There is no effective way to cure PD. Drug therapy and surgical treatment only delay the progress of the disease. Drug therapy is the primary method, but the long-term use of drugs often weakens the curative effects and produces some side effects. The clinical diagnosis of PD is currently based on motor characteristics. When the motor characteristics of PD manifest, approximately 60-70% of the neurons are lost [6]. Because the early symptoms of PD are mild and generally considered the result of aging, the early diagnosis of PD is very difficult, and there is a certain rate of misdiagnosis. At the time of diagnosis, most patients exhibit obvious behavior disorders and accumulated pathological changes. Therefore, to provide better clinical treatment for PD patients and help researchers find new treatment methods, it is urgent to identify sensitive early biomarkers.

Biomarkers are indicators that objectively reflect normal physiological and pathological processes [7]. The most studied biomarkers are present in blood and cerebrospinal fluid. Because there is no barrier between the cerebrospinal fluid and the brain, cerebrospinal fluid is considered an ideal choice for the discovery of biomarkers of neurodegenerative diseases. Cerebrospinal fluid is more stable than blood due to the blood-brain barrier, and it is more difficult to obtain. early small changes during disease development are likely excluded in blood and cerebrospinal fluid due to the steady-state regulation of the internal environment before the critical point of decompensation is reached, but urine lacks a steady-state mechanism. Therefore, urine accommodates more changes than the relatively stable cerebrospinal fluid and blood, and it is more likely to contain sensitive early markers [8, 9]. Urine sensitively reflects the development of neurodegenerative diseases. For example, 29 differential proteins were identified in urine when there were no pathological β amyloid plaques in Alzheimer’s disease transgenic mice [10], and 18 significantly altered metabolites were identified in the urine of patients with PD and linked to disease stage [11].

The use of animal models avoids the influence of age, diet, physiological conditions and other factors on urine [12], and it may enable monitoring of the early course of disease. The overexpression of human α-synuclein A53T mutations causes two typical neuropathological abnormalities of PD, namely, the progressive loss of nigrostriatal dopaminergic neurons and the formation of α-synuclein inclusion bodies [13]. This mutation model is widely used in research related to PD [14-16]. Therefore, to examine the influence of the occurrence and development of PD on the urine proteome, urine samples of α-synuclein transgenic mice were collected at 2, 4, 6, 8 and 10 months. The urine proteome was analyzed using liquid chromatography-tandem mass spectrometry (LC-MS/MS) to identify differential proteins and provide some clues for the early diagnosis of PD.

## 1 Materials and methods

### 1.1 Experimental animals

Four-week-old clean male α-synuclein (A53T)-positive transgenic mice [C57BL/6J-Tg (PDGF-α-Synuclein^A53T^)ILAS] (n=4) were purchased from the Institute of Laboratory Animal Science Chinese Academy of Medical Sciences. The animal license is SCXK (Beijing) 2014-0011. The Institute of Basic Medical Sciences Animal Ethics Committee, Peking Union Medical College approved the animal procedures (ID: ACUC-A02-2014-008). All animals were housed in a standard environment with a 12-h light-dark cycle under controlled indoor temperature (22 ± 2°C) and humidity (65-70%).

### 1.2 Rotarod test

Mouse motor coordination was evaluated using a rotarod apparatus (developed by Institute of Materia Medica, Chinese Academy of Medical Science). The rotating speeds were 25 rpm, 30 rpm, and 35 rpm at a constant interval of 30 min and a time limit of 2 min. The fall latency was recorded. The time at which the mouse first fell off the rod was recorded as the fall latency to indicate its motor coordination ability.

Before each experiment, the mice were placed in a kinematics laboratory environment for more than 30 min to avoid short-term effects on the mice due to handling and changes in lighting, and the mice were trained at 15 rpm for 1 min to adapt to the rod spinning machine. Behavioral experiments were performed every three weeks.

### 1.3 Immunohistochemistry

Immunohistochemistry was performed after the last behavioral test. Male C57BL/6 mice of the same age were used as controls (n = 3). Tyrosine hydroxylase (TH) was detected.

After anesthesia, the mice were perfused with normal saline followed by 4% paraformaldehyde. The separated brain was fixed in 4% paraformaldehyde for 24 h, then transferred to 4% paraformaldehyde containing 30% sucrose for dehydration until the brain tissue sank to the bottom. The brain tissue was removed, embedded, dehydrated and sectioned at a thickness of 4 μm. After dewaxing in water, the sections were microwave repaired in EDTA buffer, washed with PBS buffer, placed in a 3% hydrogen peroxide solution, and incubated at room temperature for 10 min. The tissue was incubated with TH antibody (ab1823, Abcam, MA, USA) (1:100) at 4°C overnight, and a secondary antibody was added for 50 min. Freshly prepared DAB solution was added for color development, and purified water was added to terminate the color development reaction. Hematoxylin was used for counterstaining, and 1% hydrochloric acid alcohol was added for 1 s for differentiation. The slices were rinsed with tap water, and ammonia water was added to turn back to blue followed by a rinse with running water. The sections were subjected to an alcohol gradient (70-100%) for 1 min, dehydrated and dried in transparent xylene, and sealed with neutral gum. The number of TH-positive cells in the substantia nigra region of the mouse was recorded.

Three researchers who were unfamiliar with the experimental design performed all evaluations.

### 1.4 Urine collection and sample preparation

The urine samples of 2-, 4-, 6-, 8- and 10-month-old mice were collected. The urine samples collected in month 2 were used as the control group. The mice were placed in a metabolic cage alone overnight (12 h) to collect urine samples. During urine collection, no food was provided, but free drinking water was available to avoid urine pollution.

The collected urine was centrifuged at 4°C and 3000 × g for 10 min to remove cells and pellets. The supernatant was stored at −80°C. Prior to urine protein extraction, the urine samples were centrifuged at 4°C and 12000 × g for 10 min to remove cell debris. Four volumes of precooled ethanol were used, and the supernatant was precipitated at 4°C for 12 h. The above samples were centrifuged at 4°C and 12000 × g for 10 min. The pellet was resuspended in lysate (8 mol/L urea, 2 mol/L thiourea, 25 mmol/L DTT and 50 mmol/L Tris). The Bradford method was used to determine the protein concentration.

Filter-aided sample preparation (FASP)[17] was used for membrane-assisted enzymolysis of urine proteins. Protein (100 μg) was loaded onto 10-kD cutoff filter devices (Pall, Port Washington, NY) and washed sequentially with UA (8 mol/L urea, 0.1 mol/L Tris-HCl, pH 8.5) and 50 mmol/L NH4HCO3. DTT (20 mmol/L) was added to reduce proteins (37°C, 1 h), and 50 mmol/L IAA was reacted in the dark for 30 min to alkylate proteins. The proteins were digested with trypsin (Trypsin Gold, Promega, Fitchburg, WI, USA) (enzyme-to-protein ratio of 1:50) for 14 h at 37°C. The peptides were collected, desalted on Oasis HLB cartridges (Waters, Milford, MA, USA), dried via vacuum evaporation (Thermo Fisher Scientific, Bremen, Germany) and stored at −80°C.

Pooled peptides were fractionated using a high-pH reversed-phase peptide fractionation kit (84868, Thermo Fisher, USA) according to the manufacturer’s instructions. The peptide samples were eluted with a gradient of increasing acetonitrile concentrations. Ten fractions were collected, including a flow-through fraction, a wash fraction and 8 step-gradient sample fractions. Fractionated samples were dried completely and resuspended in 20 μL of 0.1% formic acid.

### 1.5 LC-MS/MS analysis

#### 1.5.1 DDA-MS

LC-MS/MS data acquisition was performed on a Fusion Lumos mass spectrometer (Thermo Scientific, Germany) coupled with an EASY-nLC 1200 high-performance liquid chromatography system (Thermo Scientific, Germany). For DDA-MS and DIA-MS modes, the same LC settings were used for retention time stability. The digested peptides were dissolved in 0.1% formic acid and loaded onto a trap column (75 µm × 2 cm, 3 µm, C18, 100 A°), and the eluent was transferred to a reversed-phase analysis column (50 µm × 250 mm, 2 µm, C18, 100 A°) with an elution gradient of 5-30% phase B (79.9% acetonitrile, 0.1% formic acid, flow rate of 0.3 μL/min) for 90 min. To enable fully automated and sensitive signal processing, the calibration kit (iRT kit, Biognosys, Switzerland) was spiked at a concentration of 1:20 v/v in all samples_◯_

For the generation of the spectral library, 10 peptide fractions were analyzed in DDA-MS mode. The parameters were set as follows. The full scan was acquired from 350 to 1 550 m/z at 60 000, and the cycle time was set to 3 s (top speed mode). The auto gain control (AGC) was set to 1e6, and the maximum injection time was set to 50 ms. The MS/MS scans were acquired in the Orbitrap at a resolution of 15000 with an isolation window of 2 Da and collision energy at 32% (HCD). The AGC target was set to 5e4, and the maximum injection time was 30 ms.

#### 1.5.2 DIA-MS

For the DIA-MS method, 20 individual samples were analyzed in DIA mode. The parameters were set as follows. The full scan was acquired from 350 to 1 550 m/z at 60000, followed by DIA scans with a resolution of 30 000, HCD collision energy of 32%, AGC target of 1e6 and maximal injection time of 50 ms. Thirty-six variable isolation windows were developed, and the calculation method of the windows was performed as follows. The DDA search results were sorted according to the number of identified peptide segments by m/z and divided into 36 groups. The m/z range of each group was the window width for collecting DIA data.

### 1.6 Data analysis

#### 1.6.1 Rotarod test and immunohistochemistry

Results are presented as the mean±SD. The data were analyzed using a t-test. The level of significance was set at 0.05.

#### 1.6.2 Urinary proteomics

To generate the spectral library, the raw data files from 10 fractions acquired in DDA mode were processed using Proteome Discoverer (version 2.3, Thermo Scientific, Germany) with the SwissProt mouse database (released in May 2019, containing 17016 sequences) appended with the iRT peptides sequences The search parameters consisted of the parent ion mass tolerance (10 ppm), fragment ion mass tolerance (0.02 Da), fixed modifications, carbamidomethylated cysteine (+58.00 Da), variable modifications, and oxidized methionine (+15.995 Da). Other settings were the default parameters. The applied false discovery rate (FDR) cutoff was 0.01 at the protein level. The results were imported to Spectronaut™ Pulsar (Biognosys, Switzerland) software to generate the spectral library of 665 proteins and 29028 fragments.

The raw DIA-MS files were imported to Spectronaut™ Pulsar with the default settings. Quantitative analyses were based on the peak areas of all fragment ions for MS2. Each protein was identified with at least two specific peptides and Q value < 0.01. The screening criteria for differential proteins were as follows: fold change of protein abundance > 2 and p value < 0.01 by the t-test.

### 1.7 Functional enrichment analysis

The online database DAVID 6.8 (https://david.ncifcrf.gov/) was used to perform the functional annotation of the differential proteins, including molecular function, cell component and biological process. Pathway analysis of differential proteins was performed using IPA software (Ingenuity Systems, Mountain View, CA, USA).

## 2 Results and Discussion

### 2.1 Rotarod test

The results showed a significant difference in months 5.5 and 6 compared to month 2. However, there was no behavioral difference between months 9 and 10.5. It is speculated that the number of experimental animals was small, or the experimental animals learned behaviors during the behavioral experiments performed every three weeks (Table 1).

**Table 1.**
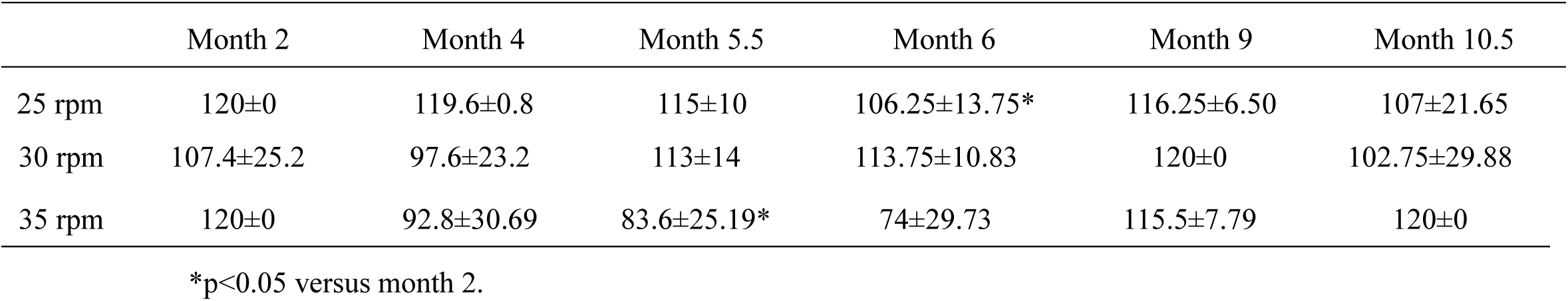
Fall latency at different time points.

|  | Month 2 | Month 4 | Month 5.5 | Month 6 | Month 9 | Month 10.5 |
| --- | --- | --- | --- | --- | --- | --- |
| 25 rpm | 120±0 | 119.6±0.8 | 115±10 | 106.25±13.75* | 116.25±6.50 | 107±21.65 |
| 30 rpm | 107.4±25.2 | 97.6±23.2 | 113±14 | 113.75±10.83 | 120±0 | 102.75±29.88 |
| 35 rpm | 120±0 | 92.8±30.69 | 83.6±25.19* | 74±29.73 | 115.5±7.79 | 120±0 |

\*p<0.05 versus month 2.

### 2.2 Immunohistochemistry

TH is the rate-limiting enzyme for dopamine synthesis. α-synuclein inclusion bodies are generally co-localized with TH. Therefore, the number of TH-positive cells reflects the loss of dopaminergic neurons. Immunohistochemistry showed a general decreasing trend in the number of TH-positive cells in the experimental group, i.e., a loss of dopaminergic neurons. There was no significant difference, which may be related to individual differences and the small number of samples (Figure 1).

**Figure 1.**
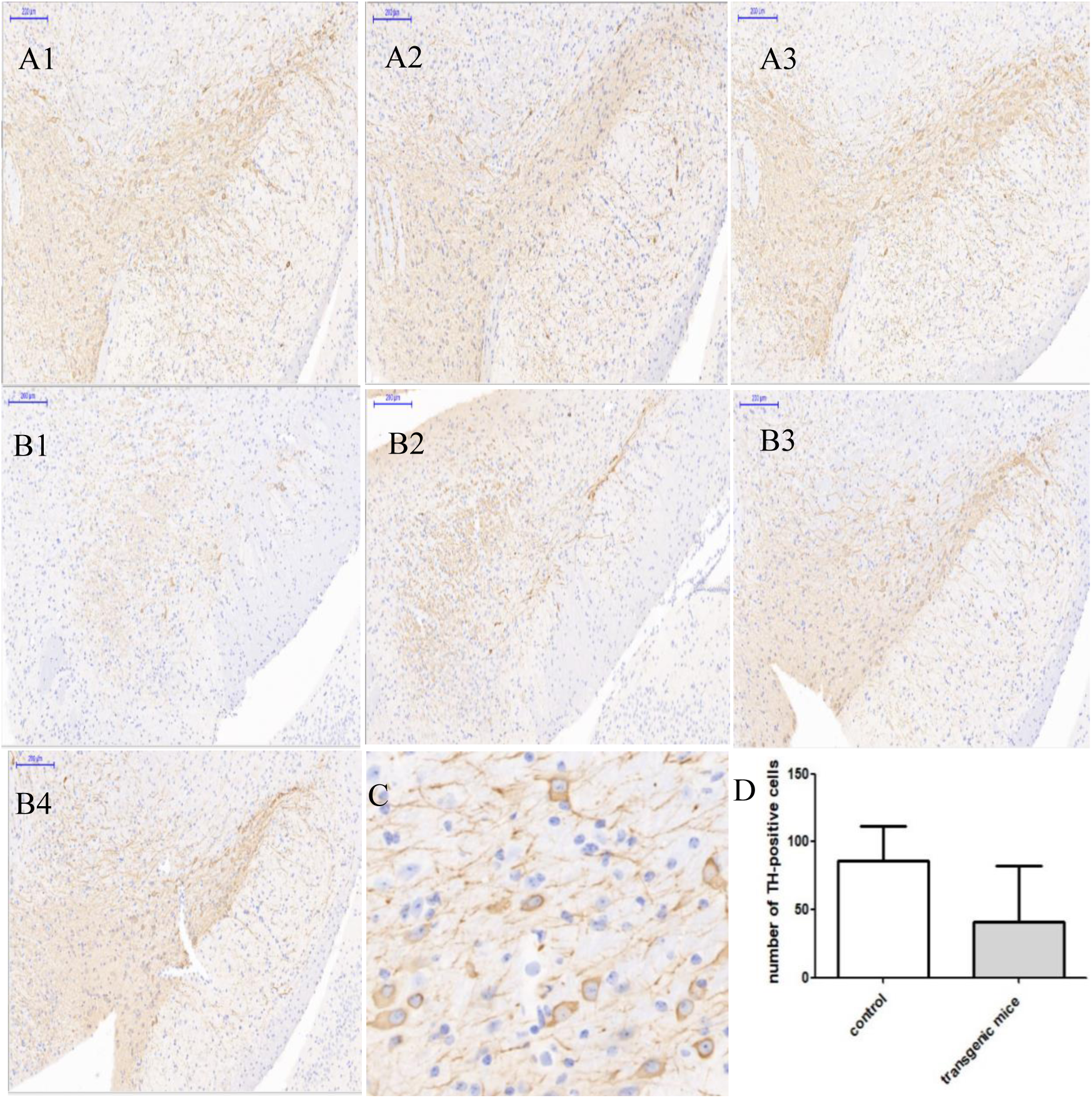
TH immunohistochemistry results. A. Control group 1-3 B. Experimental group 1-4 C. TH-positive cells D. Statistics of TH-positive cells

### 2.3 Urine proteome changes

Randomly grouped samples at each time point according to different combinations of fold changes (1.5 or 2) and p values (0.01 or 0.05) were used to calculate the average number of differential proteins under different screening conditions. This value was compared with the correct number of differential proteins under this screening condition, and the ratio approximated a false positive rate. The lower the false positive rate, the higher the reliability. Because the true positive differential proteins were also included in the results of all random groupings, the ratio of the approximate false positive rate should be slightly higher than the actual false positive rate. The condition with the lowest ratio was selected to screen for differential proteins in subsequent analyses (Table 2).

**Table 2.** Random grouping results of α-synuclein transgenic mice under different combinations of p values and fold changes.

|  | Month 4 | Month 6 | Month 8 | Month 10 |
| --- | --- | --- | --- | --- |
| 1.5, 0.05 | 15.2/9, 1.69 | 12.4/29, 0.43 | 13.8/53, 0.26 | 12.7/52, 0.24 |
| 2, 0.05 | 6.8/6, 1.13 | 5.4/15, 0.36 | 6.2/26, 0.24 | 5.7/24, 0.24 |
| 1.5, 0.01 | 2.9/3, 0.97 | 2.5/10, 0.25 | 2.7/17, 0.16 | 2.9/19, 0.15 |
| 2, 0.01 | 1.5/2, 0.75 | 1.2/8, 0.15 | 1.2/10, 0.12 | 1.1/12, 0.09 |

Note: expressed in the form of "average differential protein number/due differential protein number, ratio of the two"

Based on the random grouping, we selected the following screening criteria for differential proteins: fold change > 2 and p value < 0.01. The UniProt database was used to match the human homologous proteins that corresponded to these proteins.

DIA quantitative analysis identified 513 proteins. Seventeen human homologous differential proteins were screened in comparison to month 2. There were 2 differential proteins in month 4, of which 2 were upregulated, and 2 were downregulated. There were 7 differential proteins in month 6, of which 4 were upregulated, and 3 were downregulated. There were 5 differential proteins in month 8, of which 1 was upregulated, and 4 were downregulated. There were 10 differential proteins in month 10, of which 5 were upregulated, and 5 were downregulated. There was no continuously changed protein (Figure 2, Table 3).

**Table 3.**
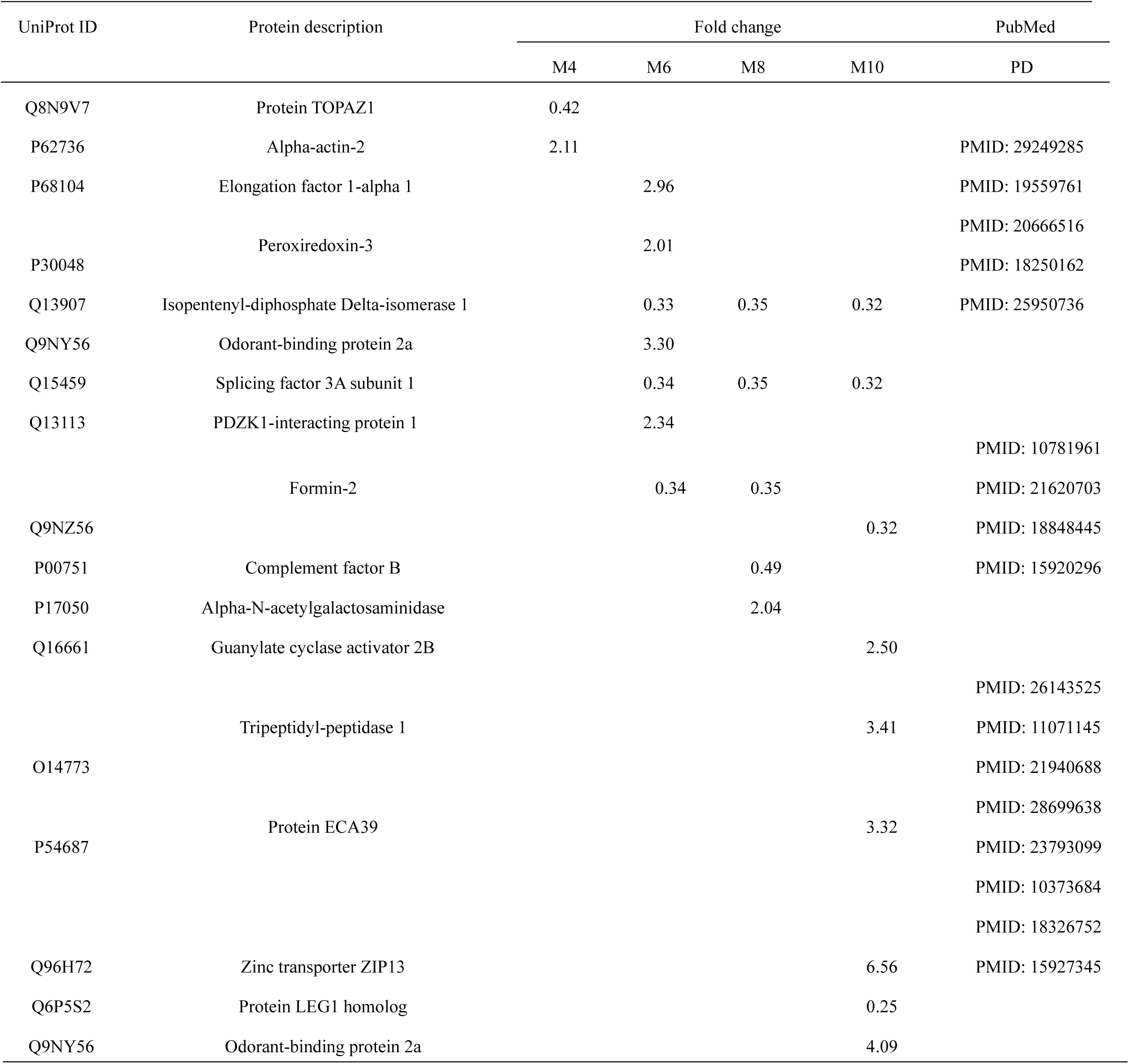
Differential proteins in α-synuclein transgenic mice.

**Figure 2.**
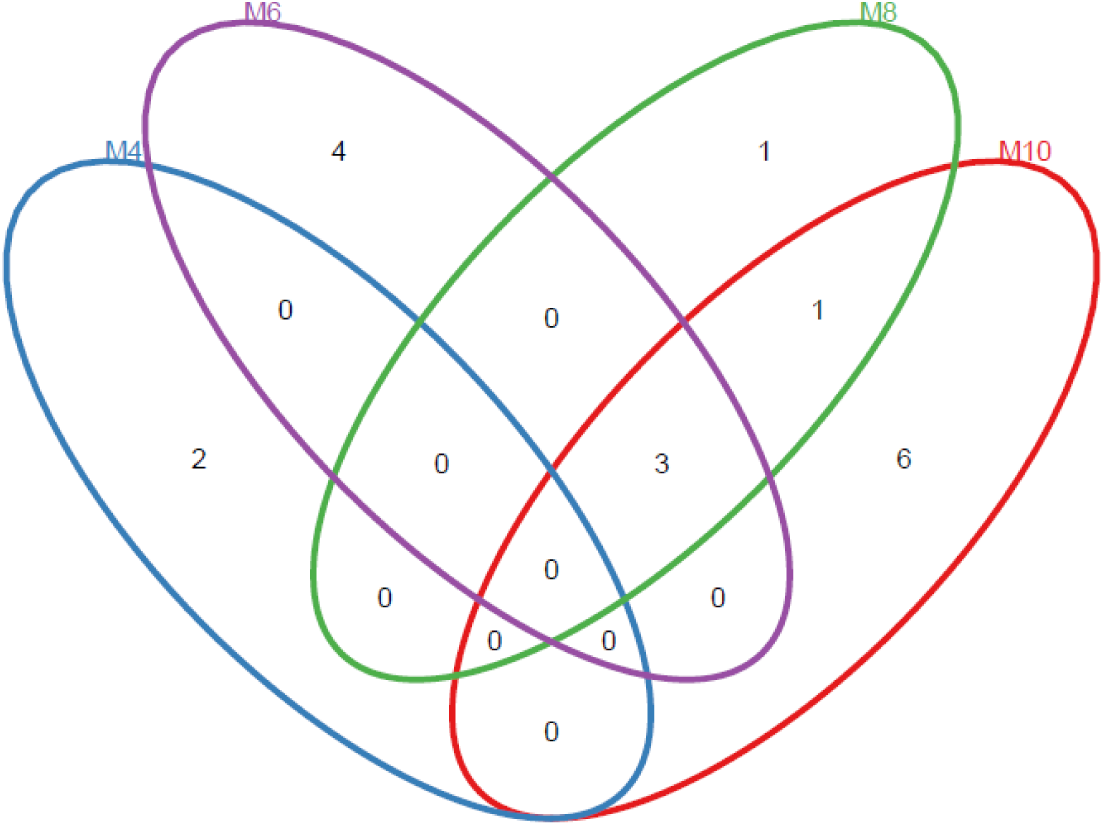
Venn diagram of differential proteins at different stages in α-synuclein transgenic mice.

There were unique differential proteins in α-synuclein transgenic mice at different time points, and there were 3 kinds of continuously changing proteins in months 6, 8 and 10, which suggests that the urine proteome has great potential to reflect the disease process in α-synuclein transgenic mice. The following specific analyses were used.

There were 2 differential proteins in month 4 prior to any significant differences in the behavior and pathology. One protein was related to PD. Actin in aortic smooth muscle is involved in various types of cell movements. The interaction between α-synuclein and hemagglutinin causes pathological changes in the actin cytoskeleton, which was verified in a previous study of α-synuclein transgenic mice [18].

Dyskinesias appeared in month 6, and 7 differential proteins were significantly changed. Four of these proteins were directly or indirectly related to PD, and 3 proteins were changed continuously in months 6, 8 and 10, Formin-2, Splicing factor 3A subunit 1 and Isopentenyl-diphosphate delta-isomerase 1. (1) Formin-2 is an actin-binding protein that is involved in the assembly and reorganization of the actin cytoskeleton [19, 20], and it is highly expressed in the developing adult central nervous system [21]. Some studies found that it was downregulated in the brain tissue of patients with Alzheimer’s disease [22]. (2) Isopentenyl-diphosphate delta-isomerase 1 is one of two isoforms (IDI1 and IDI2) of isopentenyl diphosphate isomerase (IDI) in humans. IDI1 and IDI2 may be related to the production of cholesterol metabolites in neurons, which results in the aggregation of α-synuclein during the formation of Lewy bodies [23]. (3) Peroxiredoxin-3 participates in antioxidant mechanisms, and it is upregulated in the striatum of 6-OHDA-induced PD animal models [24]. (4) Elongation factor 1-alpha 1 co-immunoprecipitates with human LRRK2, and autosomal dominant mutations in leucine-rich repeat kinase 2 (LRRK2) are the most common genetic cause of PD [25].

Five differential proteins were significantly changed in month 8, and 3 of these proteins were directly or indirectly related to PD. In addition to the proteins that changed continuously in months 6, 8 and 10, the concentration of complement factor B isoform in the cerebrospinal fluid of patients with PD was lower than that in normal subjects. The complement protein isoform in cerebrospinal fluid may be a biomarker of neurodegenerative diseases [26].

Ten differential proteins were significantly changed in month 10, and 5 were directly or indirectly related to PD. Other proteins were altered in addition to the proteins that changed continuously in months 6, 8 and 10. (1) Zinc transporter ZIP13 was altered. Zinc transporter dysfunction is related to the homeostasis of zinc ions in PD [27],and changes in zinc transporter expression and function may contribute to the development of neurodegenerative diseases[28]. The changes in Zn^2+^ content in cells may play a role in the neuronal dysfunction related to brain aging [29]. (2) Tripeptidyl-peptidase 1 was altered, and 5 patients with a mutation in the tripeptidyl-peptidase 1 gene developed PD characteristics [30-32]. (3) Protein ECA39 is cytosolic and regulates macrophage function, and it has therapeutic significance in inflammatory diseases [33]. Chronic and acute central nervous system inflammation is the cause of neurological damage [34]. PD is related to oxidative stress-mediated mitochondrial dysfunction, and the branched-chain amino acid catabolism via this protein is important to promote the tricarboxylic acid cycle, i.e., there is a certain correlation between Protein ECA39 and PD [35].

Comparing the above 17 differential proteins with the aging biomarkers [36] under the condition that “fold change > 2 and p value < 0.01”, no shared differential proteins were found. The 17 differential proteins screened in this study were related to the PD simulated in α-synuclein transgenic mice.

Under the condition of “fold change > 2, p value < 0.01”, the results of random grouping showed that the differential protein ratios in months 4, 6, 8 and 10 were 75%, 15%, 12%, and 9%, respectively. Except for month 4, the differential protein ratios in the remaining months were less than 20%. The reason for the high ratio in month 4 may be the lack of significant difference in the behavior and pathology of α-synuclein transgenic mice at this time. Month 4 belongs to the early stage of disease progression, and the change in the urine proteome was indeed small. At this time, the randomness of the differential protein obtained from screening was large, and the reliability was low.

In general, the differential proteins screened under the condition of “fold change > 2, p value < 0.01” are likely more reliable and provide more potential as urine protein biomarkers for PD. To maximize the reliability of the differential proteins in the urine proteome, the number of experimental animals should be increased in future urine biomarker studies, especially research on early stage diseases or diseases with little change.

### 2.4 Functional enrichment analysis of differential proteins

Randomly grouped samples at each time point according to different fold-changes and p-values identified the screening condition with the lowest differential protein ratio. The differential proteins obtained using these conditions are likely more reliable. In general, the stricter the screening conditions, the lower the ratio and false positive rate. However, the strict screening conditions also result in fewer differential proteins and the screening out of false-negative proteins. Because there were few differential proteins, it was impossible to perform functional enrichment analysis on the identified differential proteins using DAVID 6.8 (https://david.ncifcrf.gov/). Therefore, the functional enrichment analysis of the differential proteins screened under the condition of “fold change > 1.5, p value < 0.05” were analyzed in this experiment (Table 4).

**Table 4.**
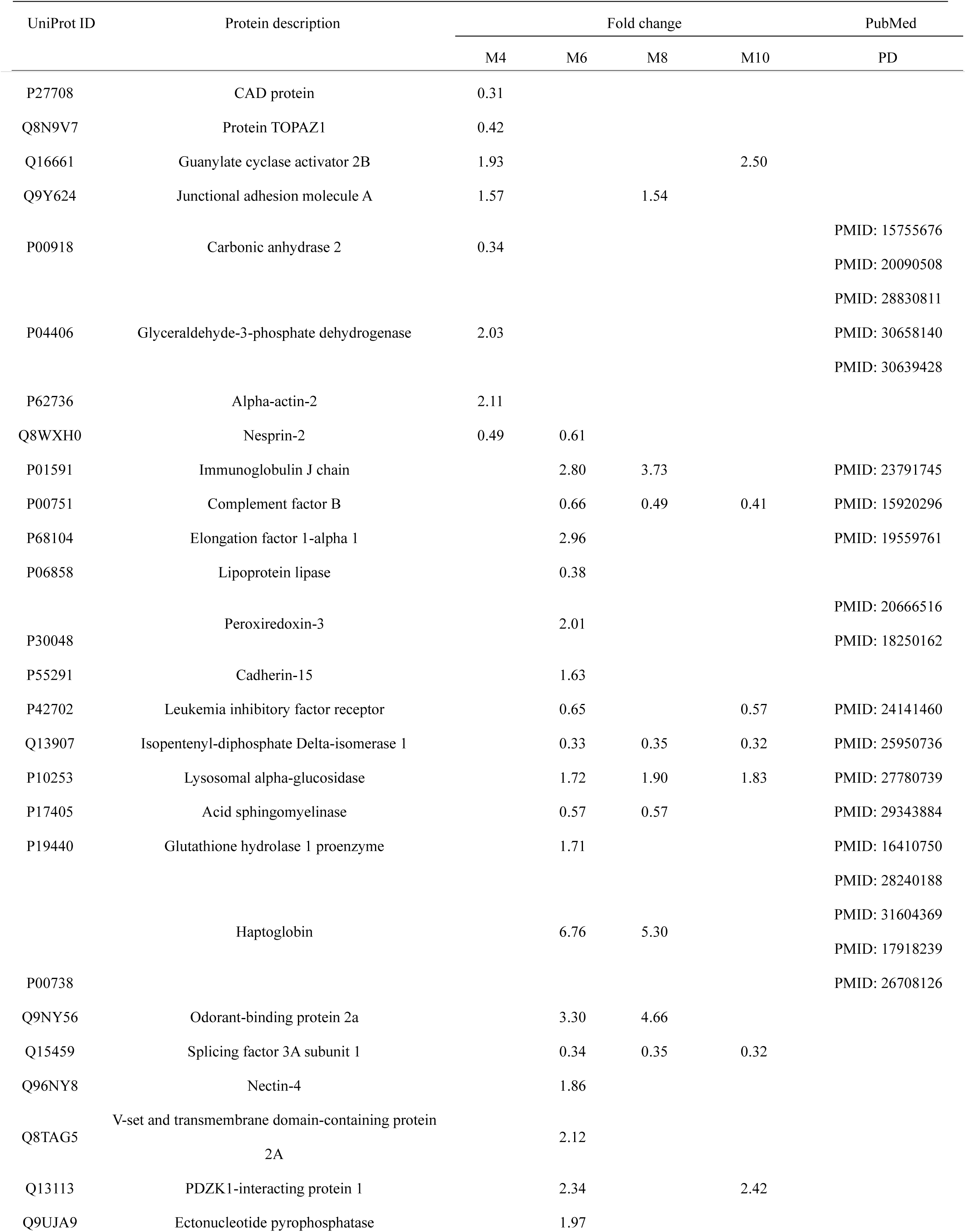
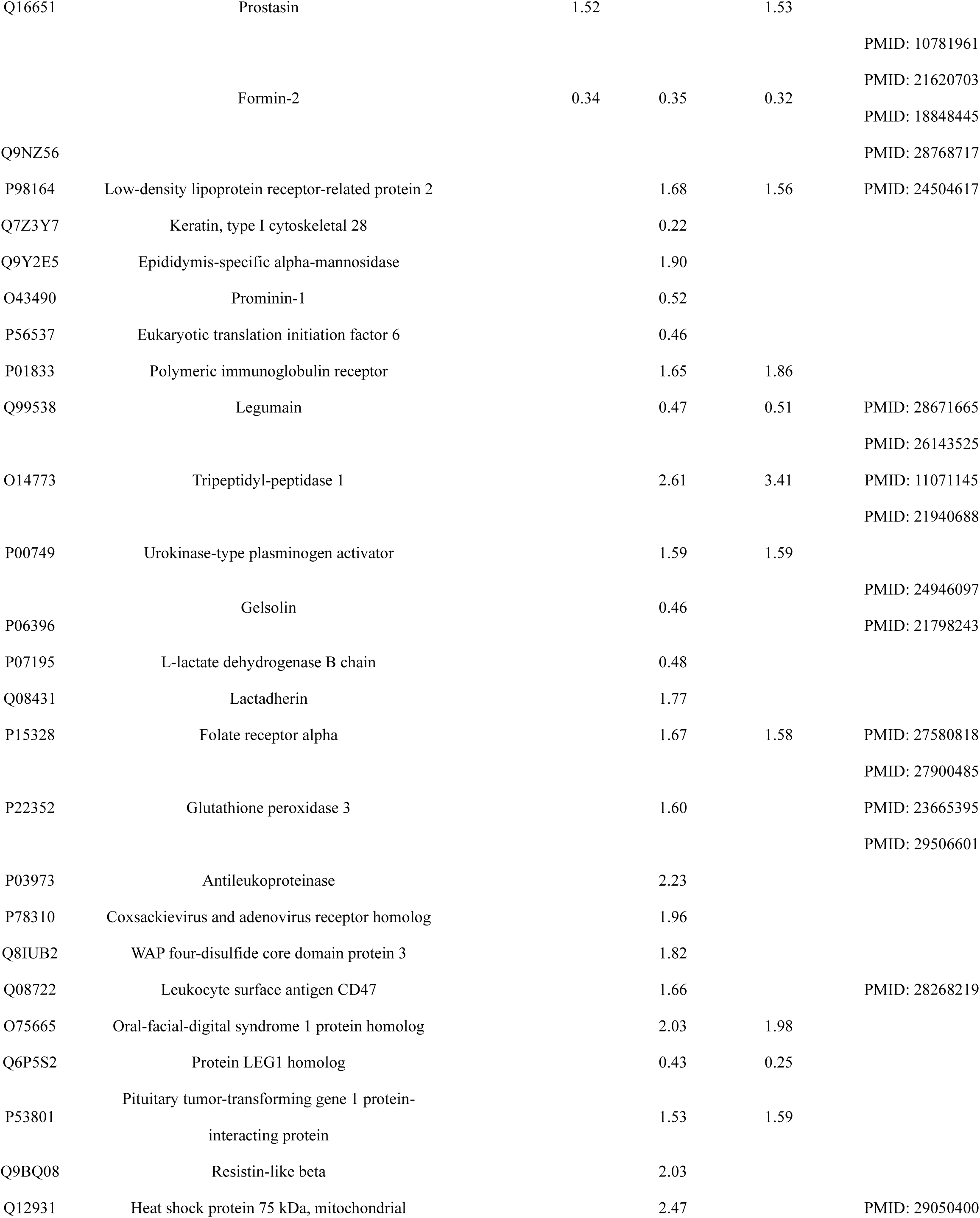
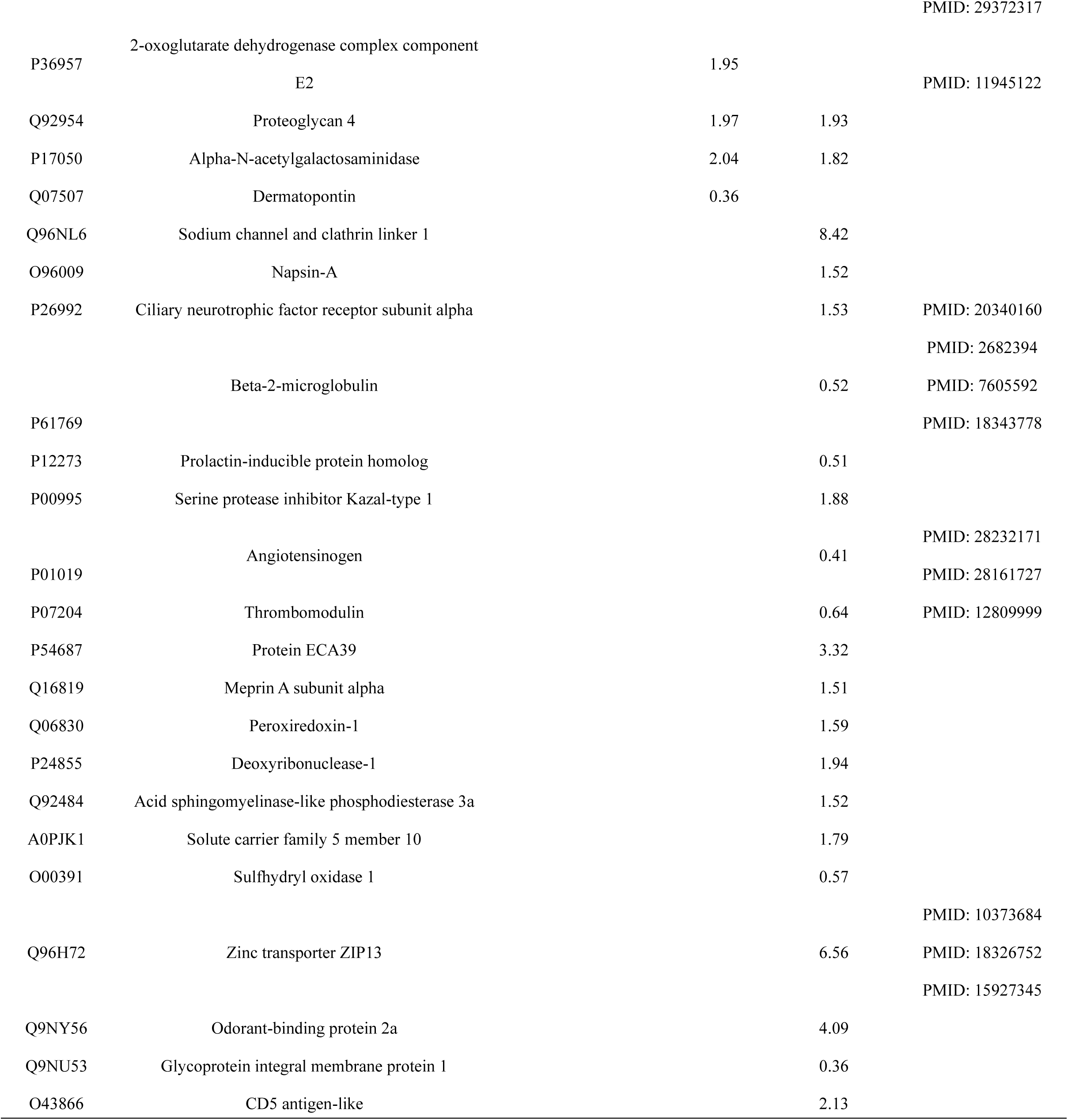
Differential proteins in α-synuclein transgenic mice (p<0.05)

**Table 5.**
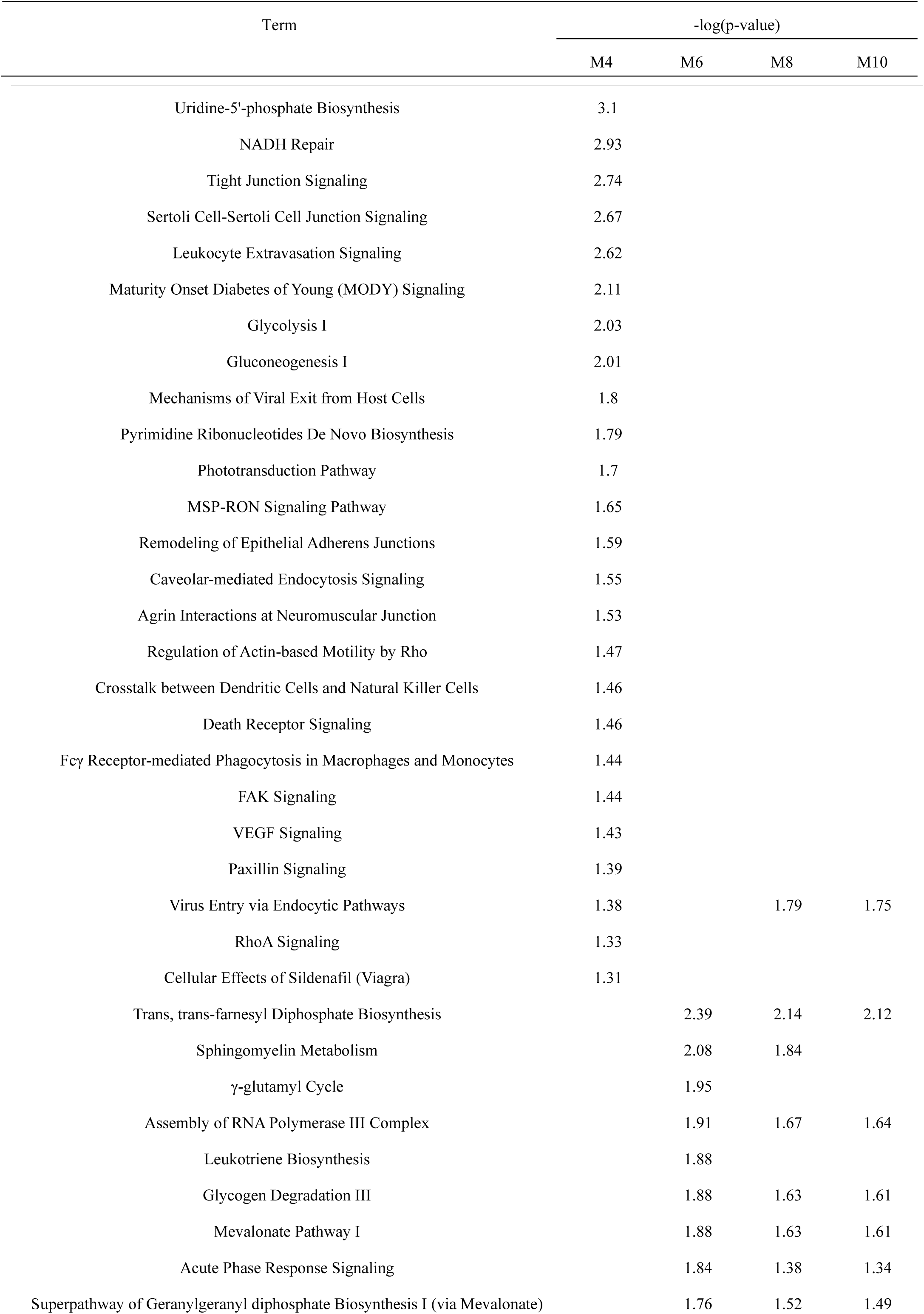
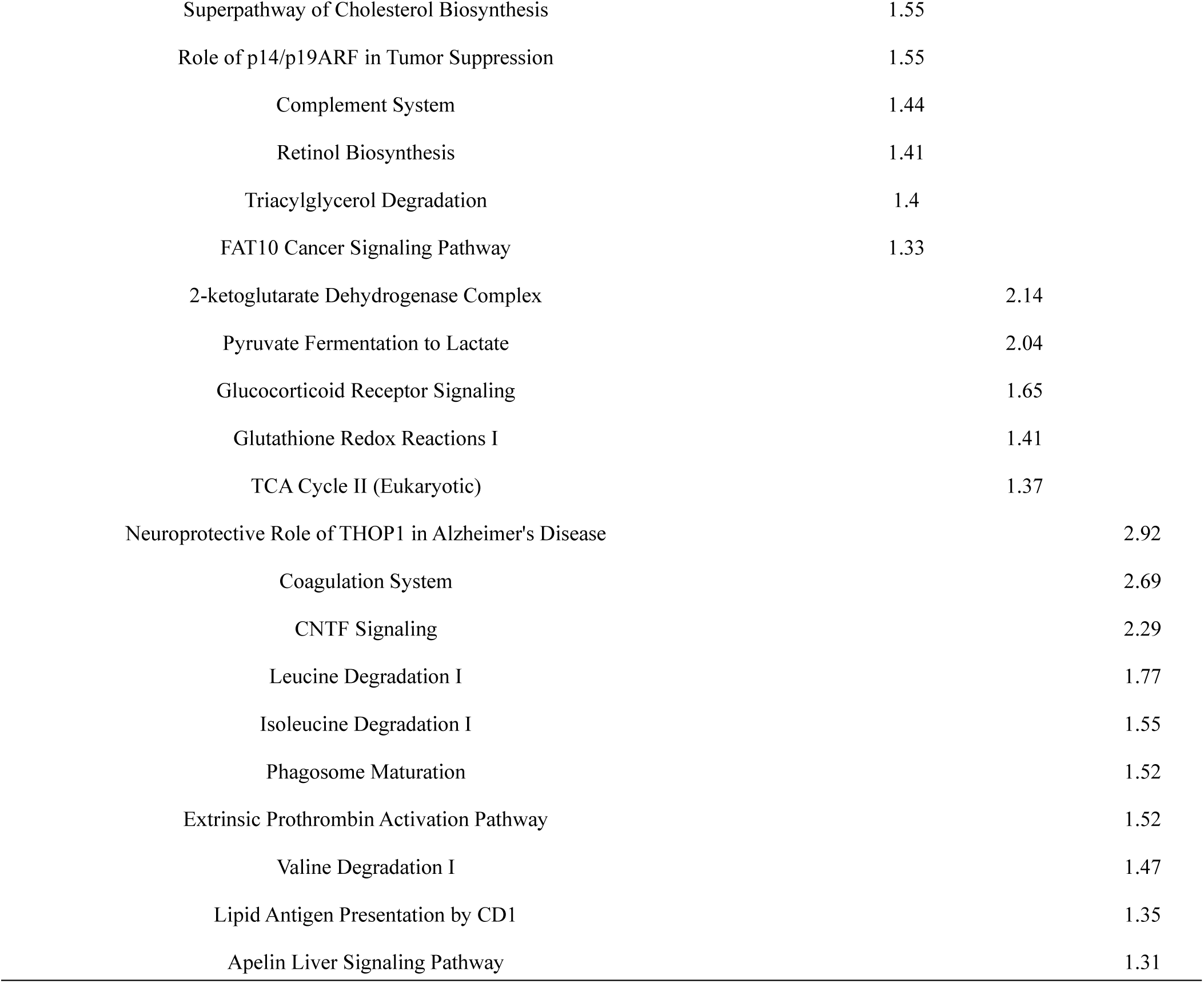
Pathway analysis of differential proteins in α-synuclein transgenic mice.

DAVID was used to annotate the function of differential proteins and show the biological process, cell composition and molecular function at each time point. IPA was used to analyze the pathways of differential proteins.

Using these screening conditions, the ratios of differential proteins in months 4, 6, 8 and 10 were 169%, 43%, 26% and 24%, respectively. The ratio of differential proteins in month 4 was greater than 100%, which primarily occurred because the time point was too early and only slightly different from the control. Therefore, the changes in urine at this time point were random, and the reliability was low. Therefore, the functional enrichment analysis in month 4 may only be used for reference.

The evaluation of biological function annotation revealed that inflammatory reaction and vitamin D metabolism began to appear in month 8, and chronic and acute central nervous system inflammation is the cause of nervous system damage in PD [34]. Vitamin D is involved in calcium and phosphate metabolism, the immune response and regulation of brain development. Vitamin D was a good candidate serum biomarker for PD and Alzheimer’s disease [37] (Figure 3A). The analysis of cell composition showed that the differential proteins were secretory proteins, mostly from the extracellular matrix and membrane (Figure 3B). The analysis of molecular function showed that most of the proteins had enzyme activity (Figure 3C). Pathway analysis showed that the pathways at different time points had unique characteristics, and some of the proteins were related to PD (Table 4). For example, inflammation occurred in month 4, such as leukocyte extravasation signaling. Tight junction signaling and endocytosis are two of the most likely pathways associated with sporadic PD [38]. Remodeling of epithelial adherens junctions and agrin interactions at the neuromuscular junction are two of the most affected pathways in PD [39]. Acute phase response signaling and various signals related to oxidative stress appeared beginning at month 6, which suggests a certain relationship between PD and immune disease.

**Figure 3A.**
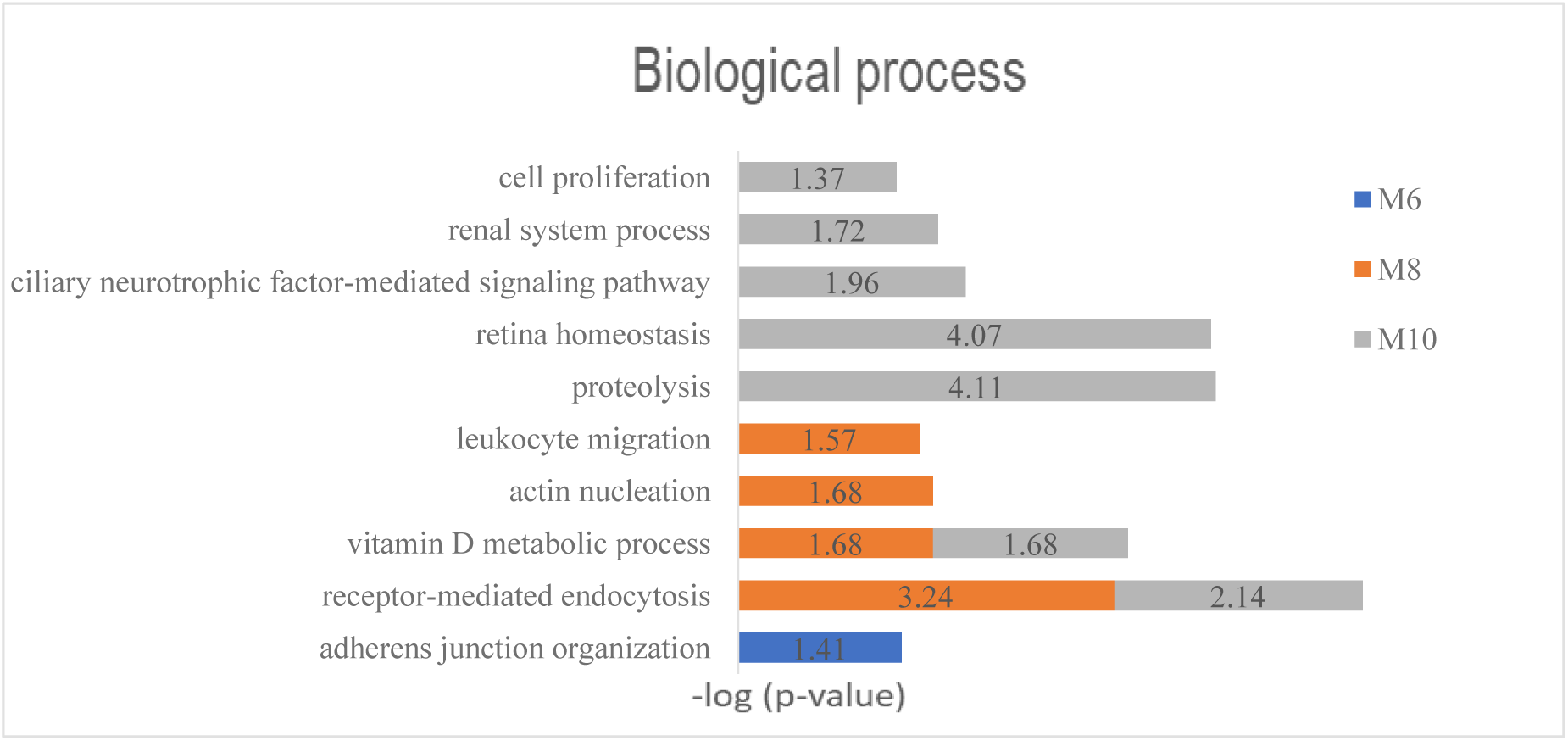
Biological process analysis of differential proteins in α-synuclein transgenic mice.

**Figure 3B.**
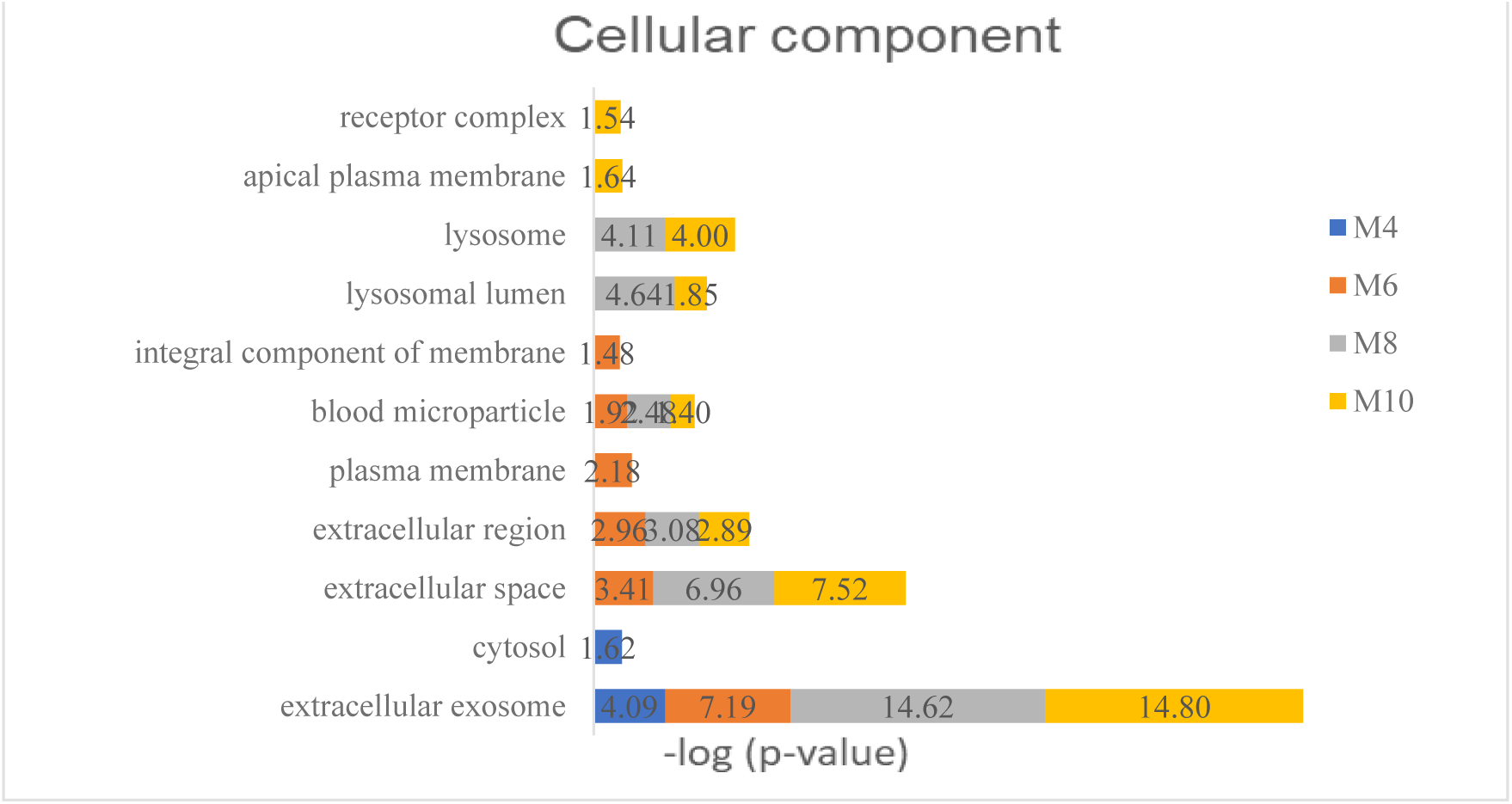
Cell component analysis of differential proteins in α-synuclein transgenic mice.

**Figure 3C.**
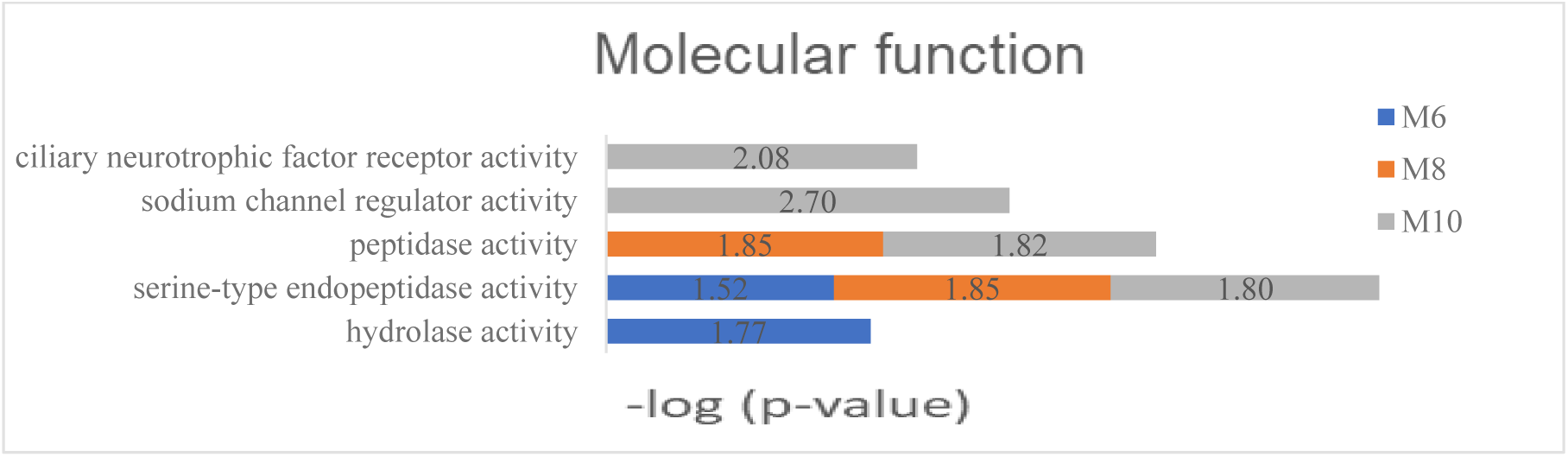
Molecular function analysis of differential proteins in α-synuclein transgenic mice.

## 3 Conclusion

The results of the present study showed that the urinary proteome reflected the disease progression changes of α-synuclein transgenic mice. There were continuous differential proteins and unique differential proteins at different time points, and more than half of the differential proteins were directly or indirectly related to PD, which suggests that urine has great potential for diagnosing and indicating the occurrence and development of PD. Urine proteome research for PD has great potential. The use of different combinations of biomarkers provides new ideas and clues for the early diagnosis of neurodegenerative diseases, further improves the reliability of diagnosis, and aids in the performance of large-scale clinical research.

## References

1. Deng, H. and L. Yuan, Genetic variants and animal models in SNCA and Parkinson disease. Ageing Research Reviews, 2014. 15: p. 161–176.

2. Yang, Y. and N. Wood, Ds, Molecular basis of Parkinson’s disease. Neuroreport, 2009. 20(2): p. 150.

3. Delenclos, M., et al., Biomarkers in Parkinson’s disease: Advances and strategies. Parkinsonism & Related Disorders, 2016. 22(s1–3): p. S106.

4. Yang, Y.X., N.W. Wood, and D.S. Latchman, Molecular basis of Parkinson’s disease. NeuroReport, 2009. 20(2): p. 150–156.

5. Delenclos, M., et al., Biomarkers in Parkinson’s disease: Advances and strategies. Parkinsonism & Related Disorders, 2016. 22: p. S106–S110.

6. Mehta, S.H. and C.H. Adler, Advances in Biomarker Research in Parkinson’s Disease. Current Neurology and Neuroscience Reports, 2015. 16(1): p. 7.

7. Strimbu, K. and J.A. Tavel, What are biomarkers? Curr Opin HIV AIDS, 2010. 5(6): p. 463–6.

8. Gao, Y., Urine—an untapped goldmine for biomarker discovery? Science China Life Sciences, 2013. 56(12): p. 1145–1146.

9. Adachi, J., et al., The human urinary proteome contains more than 1500 proteins, including a large proportion of membrane proteins. Genome biology, 2006. 7(9): p. R80–R80.

10. Zhang, F., et al., Early Candidate Urine Biomarkers for Detecting Alzheimer’s Disease Before Amyloid-beta Plaque Deposition in an APP (swe)/PSEN1dE9 Transgenic Mouse Model. J Alzheimers Dis, 2018. 66(2): p. 613–637.

11. Luan, H., et al., Comprehensive urinary metabolomic profiling and identification of potential noninvasive marker for idiopathic Parkinson’s disease. Sci Rep, 2015. 5: p. 13888.

12. Wu, J. and Y. Gao, Physiological conditions can be reflected in human urine proteome and metabolome. Expert Review of Proteomics, 2015. 12(6): p. 623–636.

13. Spillantini, M.G., et al., Alpha-synuclein in Lewy bodies. Nature, 1997. 388(6645): p. 839–40.

14. Chen, X., et al., Longitudinal Metabolomics Profiling of Parkinson’s Disease-Related alpha-Synuclein A53T Transgenic Mice. PLoS One, 2015. 10(8): p. e0136612.

15. Zhang, S., Q. Xiao, and W. Le, Olfactory dysfunction and neurotransmitter disturbance in olfactory bulb of transgenic mice expressing human A53T mutant alpha-synuclein. PLoS One, 2015. 10(3): p. e0119928.

16. Rothman, S.M., et al., Metabolic abnormalities and hypoleptinemia in alpha-synuclein A53T mutant mice. Neurobiol Aging, 2014. 35(5): p. 1153–61.

17. Wiśniewski, J.R., et al., Universal sample preparation method for proteome analysis. Nature Methods, 2009. 6(5): p. 359–362.

18. Ordonez, D.G., M.K. Lee, and M.B. Feany, alpha-synuclein Induces Mitochondrial Dysfunction through Spectrin and the Actin Cytoskeleton. Neuron, 2018. 97(1): p. 108-124.e6.

19. Pfender, S., et al., Spire-type actin nucleators cooperate with Formin-2 to drive asymmetric oocyte division. Curr Biol, 2011. 21(11): p. 955–60.

20. Azoury, J., et al., Spindle positioning in mouse oocytes relies on a dynamic meshwork of actin filaments. Curr Biol, 2008. 18(19): p. 1514–9.

21. Leader, B. and P. Leder, Formin-2, a novel formin homology protein of the cappuccino subfamily, is highly expressed in the developing and adult central nervous system. Mech Dev, 2000. 93(1-2): p. 221–31.

22. Agis-Balboa, R.C., et al., Formin 2 links neuropsychiatric phenotypes at young age to an increased risk for dementia. Embo j, 2017. 36(19): p. 2815–2828.

23. Nakamura, K., et al., Isopentenyl diphosphate isomerase, a cholesterol synthesizing enzyme, is localized in Lewy bodies. Neuropathology, 2015. 35(5): p. 432–40.

24. Lessner, G., et al., Differential proteome of the striatum from hemiparkinsonian rats displays vivid structural remodeling processes. J Proteome Res, 2010. 9(9): p. 4671–87.

25. Gillardon, F., Interaction of elongation factor 1-alpha with leucine-rich repeat kinase 2 impairs kinase activity and microtubule bundling in vitro. Neuroscience, 2009. 163(2): p. 533–9.

26. Finehout, E.J., Z. Franck, and K.H. Lee, Complement protein isoforms in CSF as possible biomarkers for neurodegenerative disease. Dis Markers, 2005. 21(2): p. 93–101.

27. Rolfs, A. and M.A. Hediger, Metal ion transporters in mammals: structure, function and pathological implications. J Physiol, 1999. 518(Pt 1): p. 1–12.

28. Leung, K.W., et al., Expression of ZnT and ZIP zinc transporters in the human RPE and their regulation by neurotrophic factors. Invest Ophthalmol Vis Sci, 2008. 49(3): p. 1221–31.

29. Mocchegiani, E., et al., Brain, aging and neurodegeneration: role of zinc ion availability. Prog Neurobiol, 2005. 75(6): p. 367–90.

30. Di Giacopo, R., et al., Protracted late infantile ceroid lipofuscinosis due to TPP1 mutations: Clinical, molecular and biochemical characterization in three sibs. J Neurol Sci, 2015. 356(1-2): p. 65–71.

31. Simonati, A., et al., A CLN2 gene nonsense mutation is associated with severe caudate atrophy and dystonia in LINCL. Neuropediatrics, 2000. 31(4): p. 199–201.

32. Le, N.M. and S. Parikh, Late infantile neuronal ceroid lipofuscinosis and dopamine deficiency. J Child Neurol, 2012. 27(2): p. 234–7.

33. Papathanassiu, A.E., et al., BCAT1 controls metabolic reprogramming in activated human macrophages and is associated with inflammatory diseases. Nat Commun, 2017. 8: p. 16040.

34. Mazzio, E.A., et al., Natural product HTP screening for attenuation of cytokine-induced neutrophil chemo attractants (CINCs) and NO2-in LPS/IFNgamma activated glioma cells. J Neuroimmunol, 2017. 302: p. 10–19.

35. Tonjes, M., et al., BCAT1 promotes cell proliferation through amino acid catabolism in gliomas carrying wild-type IDH1. Nat Med, 2013. 19(7): p. 901–908.

36. Li, X. and Y. Gao, Potential urinary aging markers of 20-month-old rats. PeerJ, 2016. 4: p. e2058.

37. Bivona, G., et al., Non-Skeletal Activities of Vitamin D: From Physiology to Brain Pathology. Medicina (Kaunas), 2019. 55(7).

38. Hu, Y., et al., A Pooling Genome-Wide Association Study Combining a Pathway Analysis for Typical Sporadic Parkinson’s Disease in the Han Population of Chinese Mainland. Mol Neurobiol, 2016. 53(7): p. 4302–18.

39. Karim, S., et al., Gene expression analysis approach to establish possible links between Parkinson’s disease, cancer and cardiovascular diseases. CNS Neurol Disord Drug Targets, 2014. 13(8): p. 1334–45.

